# Multidecadal Remote Sensing of *Macrocystis Pyrifera* in Argentina

**DOI:** 10.64898/2026.07.09.737405

**Authors:** Ashland K. Aguilar, Carolina Pantano, Henry F. Houskeeper, Tom W. Bell

**Affiliations:** Department of Applied Ocean Physics and Engineering, Woods Hole Oceanographic Institution, Woods Hole, Massachusetts, United States of America; Por El Mar Foundation, Buenos Aires, Argentina

## Abstract

The Southern Hemisphere is home to extensive forests of giant kelp (*Macrocystis pyrifera*), including in Argentina and the southern islands of Tierra del Fuego, which has been proposed as a potential climate refugium. This study presents the first regional time series of *M. pyrifera* canopy dynamics in Argentina using Landsat satellite imagery from 1985 to 2023. The forests analyzed support 247.61 km² of emergent canopy and are situated in the coastal waters of Argentina and a small portion of Chilean islands, with 4%, 28%, and 68% in the Chubut, Santa Cruz, and Tierra del Fuego A.e.I.A.S, respectively. The small portion of Chilean Islands are included as part of the Tierra del Fuego province analyses. Range limits were scrutinized, in part, using expert knowledge and multisatellite comparisons. Linear regression shows that between 1998 and 2023, 7.4% of kelp sites exhibited a significant trend in annual canopy area, with all observed significant trends in the positive direction. Partitioning by province boundaries, linear regression produces significant positive increases in kelp canopy area across all three provinces, although reassessment when longer temporal continuity is also warranted, where available. Observed seawater nitrate concentrations were high throughout the region (7–23 µmol L⁻¹), suggesting that nitrate availability was not a primary driver of canopy variability. However, positive relationships between kelp canopy and the Antarctic Oscillation suggest that regional climate variability—which alters sea surface temperature and other oceanographic conditions—may be exerting a strong influence on kelp dynamics in this region. These findings document relative stability of kelp forest area in Argentina over the most recent two and a half decades and provide preliminary evidence supporting possible increases in kelp area for the region.

## Introduction

Kelp forests are highly productive marine ecosystems that support high biodiversity through structural complexity and provide numerous ecosystem services [1–3]. The species that forms the foundation of this ecosystem is *Macrocystis pyrifera* or giant kelp, the largest and most widely distributed canopy-forming kelp. *M. pyrifera* can be found across coastal subtropical and temperate regions worldwide [4,5]. Kelp forest ecosystems contribute significantly to fisheries, nutrient cycling, and coastal economies [6–8]. Kelp forests are experiencing declines in several regions across the world associated with warming sea surface temperatures (SST), overgrazing, coastal development, and extreme events such as marine heatwaves [9–11]. However, kelp forest status and trends are highly variable within and among different regions [9,10]. Key environmental drivers, including SST, nutrients, turbidity, and wave energy, modify kelp survival, reproduction, and recruitment [4,12–14]. Overgrazing by herbivores, such as sea urchins, is also associated with ecosystem phase shifts, for example when urchins convert kelp forest habitats to barren rocky reef [15,16]. These phase shifts have been observed in regions along the coast of Tasmania and California, where changes in ocean currents, confounding stressors, and a fundamental behavioral change of urchins towards greater active feeding has led to widespread proliferation of urchin barrens [15,17,18]. In addition to local disturbances, large-scale climate oscillations, such as the El Niño–Southern Oscillation (ENSO), North Pacific Gyre Oscillation (NPGO), and Pacific Decadal Oscillation (PDO), can influence kelp forest dynamics regionally over interannual to decadal timescales [17–20]. Multiple confounding environmental factors can impact kelp forests simultaneously, making it difficult to tease apart key driving factors in an area or region. Despite the noted widespread declines, some remote regions may serve as climate refugia where kelp forests persist in localized areas of reduced disturbance or stress [21–23]. Understanding where kelp forests remain resilient to these pressures is critical for predicting future ecosystem responses and informing conservation strategies [24].

Kelp forests are highly dynamic, responding to both short- (e.g., seasonal nutrient fluxes, episodic storms) and long-term (e.g., decadal climate oscillations, increasing ocean temperatures) variability, with complex spatial dynamics [25,26]. Assessing kelp forest trends thus requires monitoring that is both spatially and temporally expansive. Traditional field-based surveys, such as diver observations, boat surveys, and aerial imagery, have been essential for localized ecological studies but are limited in spatial and temporal scope [27–29]. Remote sensing, specifically using the multidecadal Landsat satellite missions, supports large-scale and long-term monitoring of canopy-forming kelp forests. Briefly, satellite detection of floating kelp is based on the enhanced near-infrared reflectance of emergent vegetative material relative to that of seawater. Landsat imagery provides 30 m spatial resolution and an 8–16 day revisit time, offering continuous, global observations of kelp forest canopies at an ecologically relevant scale [17,30]. Temporal continuity of the Landsat program can, depending on region, support kelp forest monitoring from 1984 (or earlier, depending on resolution and radiometric requirements) to present. Long-term and continuous satellite datasets are particularly useful for studying remote areas where persistent environmental monitoring is most challenging. These spatial time series data are essential for disentangling decadal variability from long-term ecological change, especially in the face of a changing climate and increasing anthropogenic stressors.

*M. pyrifera* is influenced by a combination of environmental drivers that vary across both space and time. Some key environmental drivers include SST, nutrient availability (most commonly assessed using seawater nitrate concentration), and water clarity. In many regions, *M. pyrifera* has been shown to be highly sensitive to temperature stress, with elevated SST and marine heatwaves linked to reduced growth, canopy loss, and range contractions [9–11]. Nutrient limitation, especially in upwelling systems, can further constrain growth during periods of low nitrate availability [12,31] These drivers are always independent as temperature and nutrient availability frequently covary in upwelling systems, making it difficult to isolate their individual effects on kelp growth and survival [32]. Water clarity also plays a role in kelp recruitment and persistence. High turbidity can reduce light availability for photosynthesis, disrupt spore settlement by obscuring suitable substrate, and smother developing gametophytes and juveniles [13,14,33]. These effects may be particularly pronounced in areas influenced by riverine input, coastal runoff, or sedimentation from nearby urban development. In addition to local oceanographic conditions, regional kelp forest trends can be modulated by broader climate oscillations. Variability in temperature and nutrient availability is influenced, in part, by basin-scale climate oscillations such as the AAO, SAM, and ENSO. The asynchronized manifestation of these modes can produce nonlinear and spatially heterogeneous signatures across various regional or local scales [17,18,34].

Coastal waters of South America contain expansive *M. pyrifera* forests with highly variable status. For example, the Tierra del Fuego region has previously been described as a potential kelp forest refugia, i.e., a zone where locally favorable conditions such as lower levels of anthropogenic stressors and warming events results in heightened stability of forest canopy [21,34]. However, in other regions, such as waters near the more densely populated urban area of Ushuaia, Argentina, higher levels of anthropogenic effects, including high volumes of runoff, negatively impact kelp recruitment and understory macroalgae assemblage [13]. Prior studies investigating kelp forests in South America found that *M. pyrifera* forests have remained relatively stable for decades in the waters of Isla de los Estados [21] and the Falkland (Malvinas) Islands [35], and perhaps even longer than a century across the marine ecoregions of the Channels and Fjords of Southern Chile, the Falkland Islands (Malvinas), and the island of South Georgia [34]. Moreover, there is also evidence that *M. pyrifera* can adapt and expand to new areas with glacial retreat in the southern Magellan region [36]. While previous research has provided valuable information at localized scales [37,38], larger scale time-series analyses are still lacking for the coast of Argentina, and would provide better context on (e.g., latitudinal) variability in these past results.

This study presents the first regional assessment of *M. pyrifera* canopy dynamics along the coastline of Argentina spanning 1985-2023. However, more frequent continuity of the time series began in 1998, providing 25 years of near continuous Landsat satellite imagery from 1998–2023. The primary goals are to (1) document the distribution and trends of *M. pyrifera* canopy area in Argentina, which were partitioned using the boundaries of the three coastal provinces in Argentina where *M. pyrifera* occurs, (2) evaluate the influence of key relevant environmental variables on kelp canopy dynamics (SST, seawater nitrate concentration, and turbidity), and (3) investigate potential importance of the basin-scale climate oscillations AAO, SAM, and ENSO to spatiotemporal variability in kelp forests of Argentina.

## Methods

### Site Description

Coastal waters of Argentina support more than 200 macroalgal species, including canopy-forming subtidal kelps, of which *M. pyrifera* is the most prominent [5]. Floating canopies of *M. pyrifera* forests inhabit the coasts of the Chubut, Santa Cruz, and Tierra del Fuego provinces of Argentina [39], which span approximately 3,600 km of the ∼8,000 km of coastline (Fig. 1). The Navarino, Picton, Lennox, and Nueva Islands of Chile are also home to *M. pyrifera* and are treated as part of the Tierra del Fuego region for the analyses of this study. The oceanography of the region is characterized by the confluence of the warm Brazil Current and the cold Falkland/Malvinas Current (FMC) between approximately 35°S and 45°S. Circulation variability on the shelf is influenced by several factors, including decadal scale climate variability (such as the AAO), freshwater riverine discharge, and westerly winds, which fluctuate seasonally in magnitude [40]. The AAO influences coastal habitats in Argentina via the periodic strengthening and weakening of the westerly winds and associated low pressure encircling Antarctica, which can lead to changes in SST, currents, and upwelling [40–42]. These changes to the physical environment have been shown to alter biology, e.g., leading to seasonal blooms of phytoplankton [43–45] that form the base of highly productive fisheries [40].

**Fig. 1.**
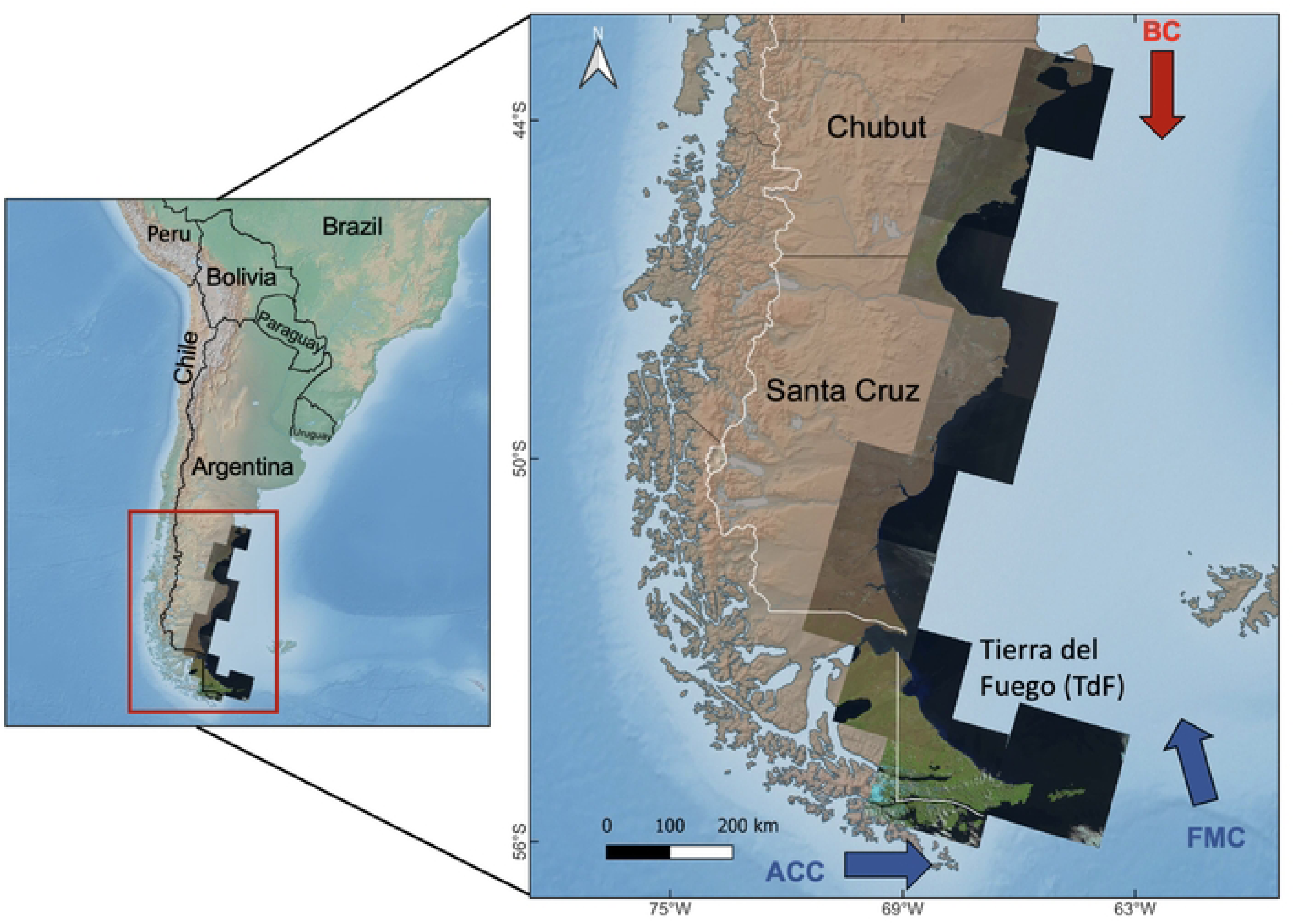
Map of the study area with the ten Landsat tiles used in the analysis overlaid along the coast. The Navarino, Picton, Lennox, and Nueva Islands of Chile are also included in the analyses as part of the Tierra del Fuego region. The red arrow is indicating the Brazil Current (BC) and the two blue arrows are indicating the Antarctic Circumpolar Current (ACC) and the Falkland/ Malvinas Current (FMC).

### Landsat Observations of Kelp Canopy Area

A time series of kelp canopy extent for coastal habitat in the Chubut, Santa Cruz, and Tierra del Fuego provinces, with the latter also including the Navarino, Picton, Lennox, and Nueva Islands of Chile, was created to assess spatiotemporal dynamics of *M. pyrifera* (Fig. 1). Imagery from 1985-2023 was accessed across four Landsat sensors: Landsat 5 Thematic Mapper, Landsat 7 Enhanced Thematic Mapper Plus, Landsat 8 Operational Land Imager, and Landsat 9 Operational Land Imager 2. Each satellite collects an image every 16 days with spatial resolution of 30 m. From 1999 through 2011, Landsat 5 and 7 operated concurrently, and after the 2013 launch of Landsat 8, image acquisition continued with overlapping missions (Landsat 7 & 8 from 2013–2021, and Landsat 8 & 9 from 2021). As a result, most years after 1999 provide imagery every 8 days. Giant kelp typically grows close to the coast (usually <1 km), and land and intertidal areas were masked using a combination of the Advanced Spaceborne Thermal Emission and Reflection Radiometer (ASTER, https://www.jspacesystems.or.jp/ersdac/GDEM/E/ ) 30 m digital elevation model and the Modified Normalized Difference Water Index (MNDWI; [29,46]). The ASTER digital elevation model masked any pixel with an elevation greater than 10 m, and MNDWI from clear images taken during a low tide were used to classify low elevation and intertidal pixels. MNDWI uses the shortwave infrared and green spectral bands to obtain an index value for each pixel, and any pixel with an MNDWI value less than 0.1 was included in the mask. The resulting land mask was validated with high resolution (∼0.5 m) satellite imagery, and misclassified intertidal areas were manually corrected in QGIS (3.32.2-Lima). The land mask was then buffered by 30 m (to account for shallow or tidally exposed substrate) and, combined with a 3 km offshore limit, applied to Landsat imagery to identify potential *M. pyrifera* habitat.

Landsat imagery (Collection 2 Level 2 Surface Reflectance) was processed in Google Earth Engine (GEE) using updates to the methods of [29,35]. Briefly, after masking for clouds and land, a globally representative random forest classifier was used to categorize pixels into kelp canopy or seawater. The classifier utilizes the blue, green, red, near infrared, and two shortwave infrared bands. The low-latitude range limit defined in the initial Landsat processing was scrutinized using a multisensor—e.g., including PlanetScope imagery with (nominally) 3 m spatial resolution shown to support monitoring of kelp canopies at fine-scale resolution [47]—comparison, and then the Landsat time series was re-processed using the updated range. The satellite datasets do not retrieve subsurface portions of *M. pyrifera* because water rapidly attenuates near-infrared light [48]. However, [49] found a significant relationship between remotely sensed *M. pyrifera* canopy and subsurface biomass (r²= 0.8976), suggesting that in many scenarios canopy area functions as a key indicator of the total *M. pyrifera* forest biomass during periods when canopy is present [29]. Finally, canopy area was estimated for each three month quarter by dividing the total number of kelp canopy presence classifications by the total number of cloud-free views for each pixel location.

### Regional Kelp Canopy Trend Analysis

Regional kelp canopy area trends were evaluated across three geographic domains, defined using the geopolitical province borders. For the northernmost domain, Chubut Province, kelp classifications were derived from the southern province boundary to the anticipated giant kelp range limit (42°S), as described by [50]. The center domain, Santa Cruz, corresponded to the province boundaries. The southernmost domain, Tierra del Fuego, encompasses the southernmost mainland coast corresponding to the province boundaries and Staten island, along with the Navarino, Picton, Lennox, and Nueva Islands of Chile. This study excludes Malivinas, South Georgia, and the South Sandwich Islands. Image classifications represent observations of emergent canopy within 30 x 30 m pixels, and classifications were then aggregated into 5 x 5 km square cells. Previous studies found that spatial synchrony of kelp canopy declines dramatically within 200 m, thus 5 x 5 km cells allow for multiple replicates in each region, while also avoiding spatial autocorrelation processes [29,51,52]. Highly sparse cells (i.e., those not containing at least 100 pixels where kelp canopy has been observed) were excluded from the analysis, which resulted in removal of only 3% of the kelp habitat from the 30 x 30 m non-aggregated dataset. If more than 25% of potential kelp habitat were obscured by cloud cover across the three-month period, the entire 5 x 5 km cell was treated as missing data for that period. An annual time series was then created for each cell using the maximum seasonal canopy area. To mitigate uncertainty due to seasonal variability, each yearly data point was required to have at least two seasons of data to be included. Because the satellite record for coastal waters of Argentina is relatively sparse between 1984 and 1998, the time period from 1998-2023 is considered in the time series analyses. All regional analyses shown in this study utilize this maximum annual canopy area time series for each 5 x 5 km cell from 1998 to 2023, but time series showing quarterly intervals and including earlier sparse data points are included in the Supplemental Materials. A generalized least squares regression model (R package nlme; [53]), was used to analyze the annual kelp canopy trends at both the province scale (summed 5 x 5 km sites within each of the three provinces) and at each of the individual 367 5 x 5 km sites with a 0, 1, or 2 year auto-regressive error structure selected by minimizing the Akaike Information Criterion (AIC). To control for multiple comparisons, p-values were corrected using the Benjamini-Hochberg false discovery rate procedure [54], with significance assessed at q < 0.05.

### SST and Seawater Nitrate Relationship Analysis

Observations of SST were obtained from NOAA’s Coral Reef Watch program (https://coralreefwatch.noaa.gov/product/5km/index_5km_sst.php ), which provides 5 km data at daily resolution. Data was downloaded spanning the period from 1985 to 2023 and covering the region between 41°S to 57°S and 58°W to 77°W. A relationship between SST and nitrate along the coast of Argentina was established using data from a previous study by [55], where water samples were collected during March-April 2005, at a depth of 9 meters across 38 locations along the Argentine Shelf, the Drake Passage, and the Northwest Antarctic Peninsula to the Bellingshausen Sea. A Generalized Additive Model (GAM) was used to analyze the relationship between temperature and seawater nitrate concentration (R package mcgv; [56]), and the relationship was then used to estimate spatial time series of seawater nitrate from the SST data. Although the nitrate and SST data are not independent and based on a relationship from a single season, this estimation is useful because it allows analysis spanning large spatial scales and temporal coverage matching that of the Landsat record. Nonetheless, potential caveats associated with the nitrate-to-SST relationship are considered in the Discussion section.

### Comparisons with Basin-Scale Climate Modes

Three oceanographic climate indices that are known to influence the study area were analyzed: the the Multivariate ENSO Index (MEI) (http://www.esrl.noaa.gov/psd/enso/mei), AAO (https://www.cpc.ncep.noaa.gov/products/precip/CWlink/daily_ao_index/aao/aao.shtml), and SAM (https://psl.noaa.gov/data/20thC_Rean/timeseries/monthly/SAM/ ). The AAO and SAM are associated with the same generalized basin-scale phenomenon but are considered separately here because the indices are constructed differently. The AAO captures strong zonal symmetry, occurs year-round, and is more active during austral spring [57]. A positive AAO phase strengthens westerly winds and shifts storm tracks poleward, enhancing upwelling and cooler conditions in sub-Antarctic regions while generally warming mid-latitude waters. Along the Patagonian shelf, this can manifest as changes in wind stress, SST, and nutrient availability that influence coastal productivity. The SAM captures north-south deviations of the westerly wind belt, which influences weather patterns in the southern hemisphere. During a positive SAM phase, the strengthening and poleward contraction of the southern westerlies is known to modify wind stress, storm track positions, and mixing regimes in the Southern Hemisphere mid-latitudes [58]. On the Patagonian shelf, several studies have linked positive SAM phases with increased SST and weakened wind-driven nutrient supply and upwelling [59,60], while shelf circulation and transport variability have also been attributed to SAM fluctuations [61]. ENSO is a recurring climate phenomenon characterized by coupled fluctuations in SST and atmospheric pressure across the equatorial Pacific. ENSO manifests as three phases: El Niño, La Niña, and a neutral state. During El Niño, trade winds in the Pacific weaken, atmospheric pressure gradients in the Pacific relax, and warm surface waters accumulate in the central and eastern Pacific. This redistribution of heat and pressure suppresses upwelling along the South American west coast and disrupts global weather patterns. In contrast, La Niña corresponds to intensified trade winds and enhanced upwelling, which cool SST in the eastern Pacific and often reinforce regional climate patterns. Although ENSO is primarily a Pacific phenomenon, it is included because of known teleconnections to other ocean regions, e.g., via disruption of global weather patterns. ENSO integrates oceanic and atmospheric anomalies in the tropical Pacific and is quantified here using MEI, with sustained positive values indicating El Niño–like conditions and sustained negative values indicating La Niña–like conditions. The relationship between variances in kelp forest canopy and climate indices were quantified using Pearson’s correlation coefficient.

### Observations of Light Attenuation and Chlorophyll a

The diffuse attenuation coefficient for downwelling irradiance (Kd), which quantifies the rate at which downward-propagating light diminishes with depth, serves as a proxy for underwater light availability and is a robust indicator of bio-optical system state [62–65]. Chlorophyll *a* (Chla) is the most ubiquitous photosynthetic pigment, and aquatic Chla concentration is commonly used to quantify phytoplankton biomass—a proxy conferring information on water clarity as well as nutrient availability. Satellite datasets of Chla plus Kd490 (Kd derived at 490 nm) were downloaded using NASA Level-3 monthly composites (at 9 km spatial resolution) from the NASA Ocean Color database for each of the SeaWiFS (1998–2010), MODIS-Aqua (2011–2019), and VIIRS-SNPP (2020–2023) missions. The three monthly datasets were then merged in MATLAB to produce continuous time series for Chla and Kd490. Kelp forest area was tested for association to Kd490 and Chla individually using Pearson’s correlation coefficient.

## Results

### Distribution of Kelp Canopy along Argentina

Dynamics of kelp forest area were assessed with a time series from 1985-2023. However, available imagery was sparse up until the launch of Landsat 7 in 1999 and, combined with data losses due to cloud cover, the early imagery time series is sparse. For consistency across regions and to minimize leverage in the time series analysis from early but sparse observations, a near-continuous time series of kelp canopy area for each region spanning 1999 – 2023 was used. However, clear imagery is available for the Chubut province during 1986 and is shown in the Supplemental Materials (Fig. S1). A time series was completed for the kelp canopy area both seasonally (Fig. S2) and annually (Fig. 2). Since kelp forest canopy dynamics are often associated with seasonal drivers, especially in Chubut and Santa Cruz [66], the annual time series was used in the statistical analyses because it was a better overall representation of the amount of kelp canopy observed year to year, i.e., it mitigated variability associated with seasonal fluctuations and timing. There were 367 sites (5 x 5 km cells) identified that had kelp forest habitat across the three provinces and met the requirement of 100 pixels of potential kelp habitat; Chubut had 34, Santa Cruz had 97, and Tierra del Fuego had 236 sites total (Fig. 3). The Santa Cruz and Tierra del Fuego provinces had approximately an order of magnitude more kelp canopy area than Chubut.

**Fig. 2.**
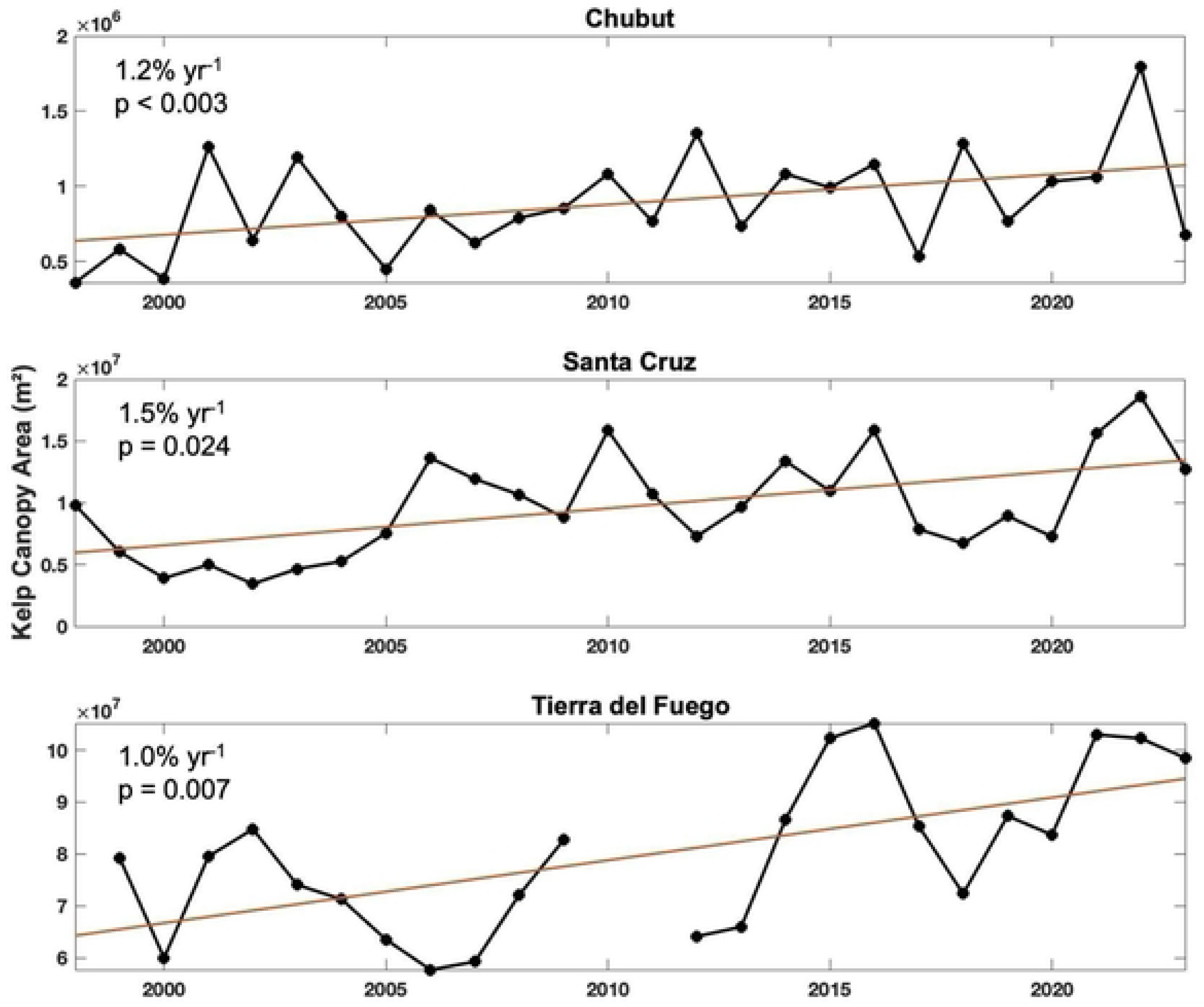
Provincial time series of annual kelp canopy area from 1998-2023.Trend line is shown in red. The annual rate of change is shown on the top left of each figure along with its associated p value.

**Fig. 3.**
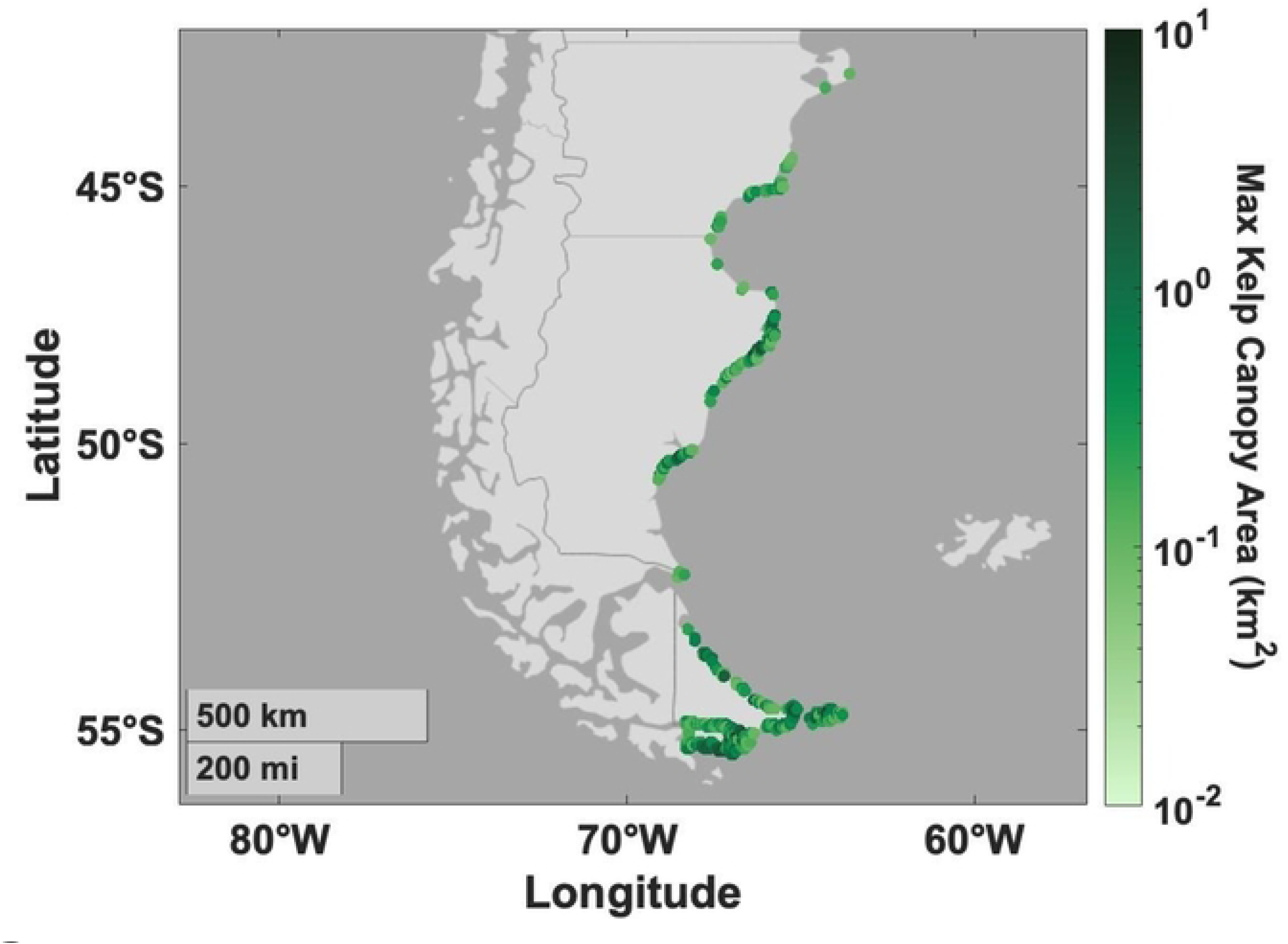
Distribution of kelp canopy from 1999-2023 displayed on a log scale. In those cells or sites, there are pixels. The color of each site corresponds to the amount of the pixels that had kelp canopy. Where sites that had high kelp canopy cover are displayed in dark green and the sites with lower kelp canopy are displayed in light green.

An investigation of the low-latitude range-limit position was performed using PlanetScope imagery, which supported extending the image-processing domain by approximately 50 km equatorward. The extension captured emergent kelp near Faro Punta Norte, which were readily classified using the Landsat method once the domain was revised (Fig. 4). The domain extension adds a small area relative to the total area analyzed here and so the added pixels are not included in the time series comparisons. Rather, the domain extension will be used to support a future reprocessing of the Argentina kelp forest Landsat record, which is hosted for public access at www.Kelpwatch.org [29].

**Fig. 4.**
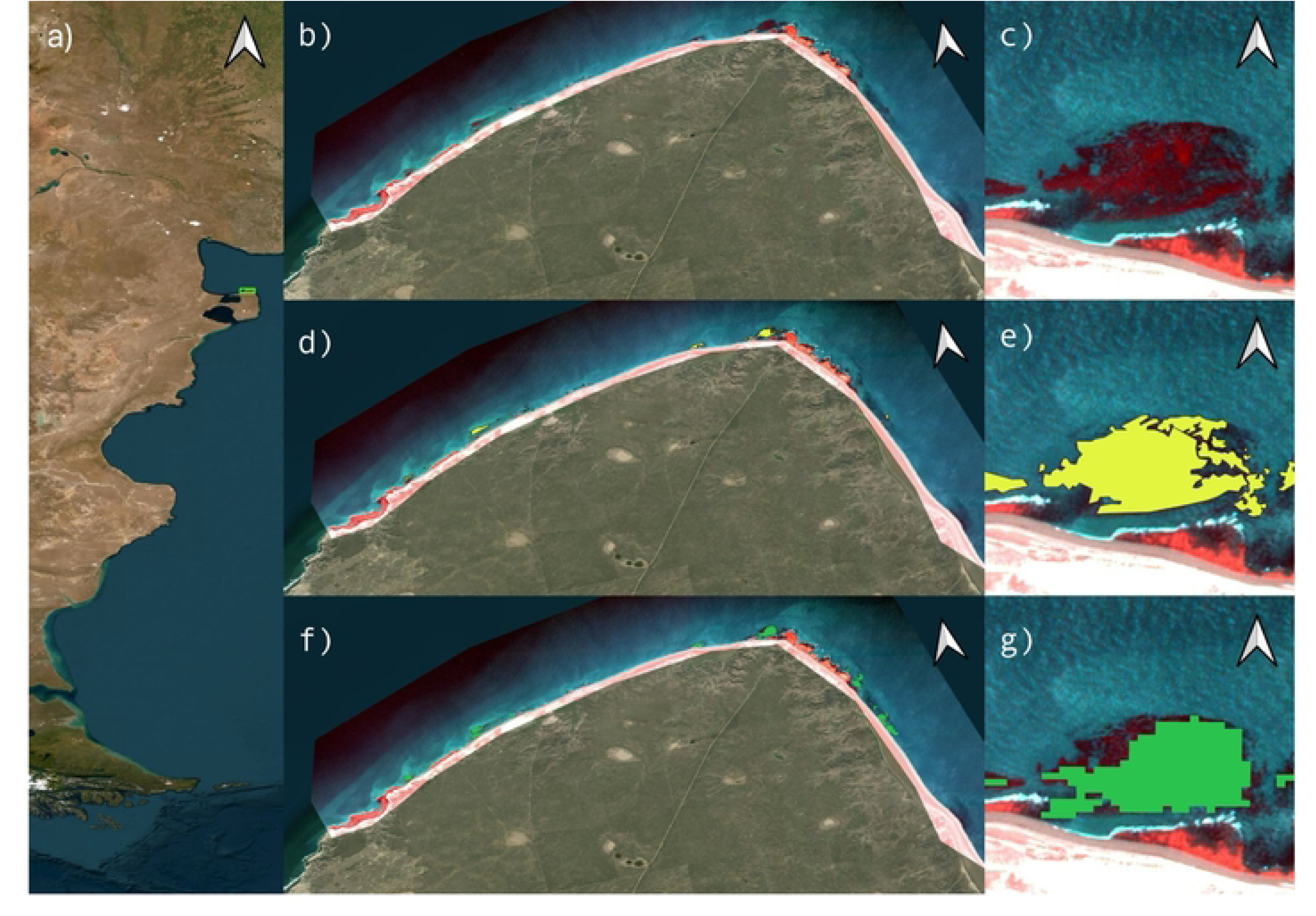
The low-latitude range limit in the Landsat record was extended northward using multisatellite comparisons including the PlanetScope products. PlanetScope pseudocolor imagery (near infrared, green, and blue) from 2022 is overlaid on a basemap in panels b throughg. A bed approximately defining the location of northernmost detection—i.e., near Faro Punta Norte, Chubut—is shown in panel c. Manual classifications of emergent kelp using a single PlanetScope scene are shown in yellow for the same regions (panels d and e), and automated classifications of emergent kelp using the Landsat record are shown in green (panels f and g).

### Change in Kelp Canopy Area across Time

A linear regression analysis indicated that annual kelp forest area increased in each of the three provinces over the period of continuous Landsat observations (1998–2023). For regional (province-scale) analyses, the results showed annualized increases at Chubut of 1.2% yr^-^¹( p = 0.003), Santa Cruz of 1.5% yr-¹ (p = 0.024), and Tierra del Fuego of 1.0% yr^-^¹ (p = 0.007), shown in Fig. 2. The cell-scale (5 x 5 km sites) analyses revealed that the regional changes were likely driven by increased area at a subset of the study domain. Following false discovery rate (FDR) correction for multiple comparisons, 27 of the 367 cells (7.4%) exhibited a significant trend, with all significant trends occurring in the positive direction. In Chubut, 2 of 34 sites (5.9%) displayed a significant positive trend. In Santa Cruz, 7 of 97 sites (7.2%) displayed a significant positive trend. In Tierra del Fuego, 18 of 236 sites (7.6%) displayed a significant positive trend. The Chubut province time series contains one early data point in 1986, followed by a multi-year data gap until 1997. Generalized least squares regression estimators can be sensitive to outliers; therefore, time series sensitivity testing was conducted by rerunning the generalized least squares regression with the early data point removed. With the initial point in 1986 included, giant kelp forest canopies are found to be increasing at a rate of 1.1% annually (p < 0.001). With the initial point removed, the observed trend in kelp canopy increased from 1.1% (p < 0.001) to 1.2% annually (p < 0.003). The results from the autocorrelation analysis are shown in figure S2.

### Relationship of Kelp Canopy to Temperature and Nitrate

SST and seawater nitrate concentration are modeled using an inverse and non-linear relationship where nitrate levels decrease as temperature increases (R²=0.936, p<0.001), shown in Fig. S3. Kelp forests in Chubut were not significantly related to SST (r= 0.0787, p = 0.6963) or seawater nitrate concentrations (r= -0.0992, p= 0.6224). Santa Cruz kelp forests were also shown to not be significantly related to SST (r= 0.1793, p= 0.3809) or nitrate concentrations (r= - 0.1916, p= 0.3484). Lastly, kelp forests in Tierra del Fuego were not strongly related to SST (r= 0.0353, p= 0.8729) or nitrate concentrations (r= 0.0043, p= 0.9846). However, differences in kelp area across provinces was consistent with generalized patterns across the regions. For example, annual nitrate concentrations were lowest in Chubut and highest in Tierra del Fuego (Fig. S4), thus annual SST was warmest in Chubut, followed by Santa Cruz, and then Tierra del Fuego.

### Relationship of Kelp Canopy Dynamics and Environment to Marine Climate Oscillations

The AAO was found to be significantly and positively related to kelp canopy dynamics in southern Argentina. Briefly, the two southern-most provinces, Santa Cruz and Tierra del Fuego, indicated r values of 0.4718 (p= 0.0150) and 0.4993 (p=0.0153), respectively, shown in (Fig. 6). Kelp canopy dynamics in the northernmost province containing *M. pyrifera*, Chubut, were not significantly related to the AAO, with r=0.3277 (p=0.0952). Despite the significant relationships identified with the AAO, the SAM was not significantly related to kelp forests in Chubut (r=0.3204, p=0.1105), Santa Cruz (r=0.3618, p=0.0693), or Tierra del Fuego (r= 0.413, p=0.0561). MEI was not strongly related to kelp forests in Chubut (r=-0.1695, p=0.3979), Santa Cruz (r=-.02052, p=0.3146) or Tierra del Fuego (r=0.1279, p=0.5609). The relationship between the AAO and SST was also analyzed, which showed a significant relationship in Chubut (r=0.4195, p=0.0087; Fig. 7) and in Santa Cruz (r=0.5002, p=0.0014; Fig. 7). The relationship between the AAO and SST was not significant in Tierra del Fuego (r=0.2409, p=0.1452; Fig. 7). The relationship between the AAO and seawater nitrate concentration was also investigated and had a significant negative relationship in Chubut (r=-0.4851, p=0.0020) and Santa Cruz (r=-0.5425, p=0.0004) but was not significantly related in Tierra del Fuego (r=-0.2370, p=0.1519). Caution is warranted when considering the relationship between AAO and seawater nitrate because the nitrate estimation is obtained using SST and uncertainties are poorly understood, for example, as a function of season or latitude.

**Fig. 5.**
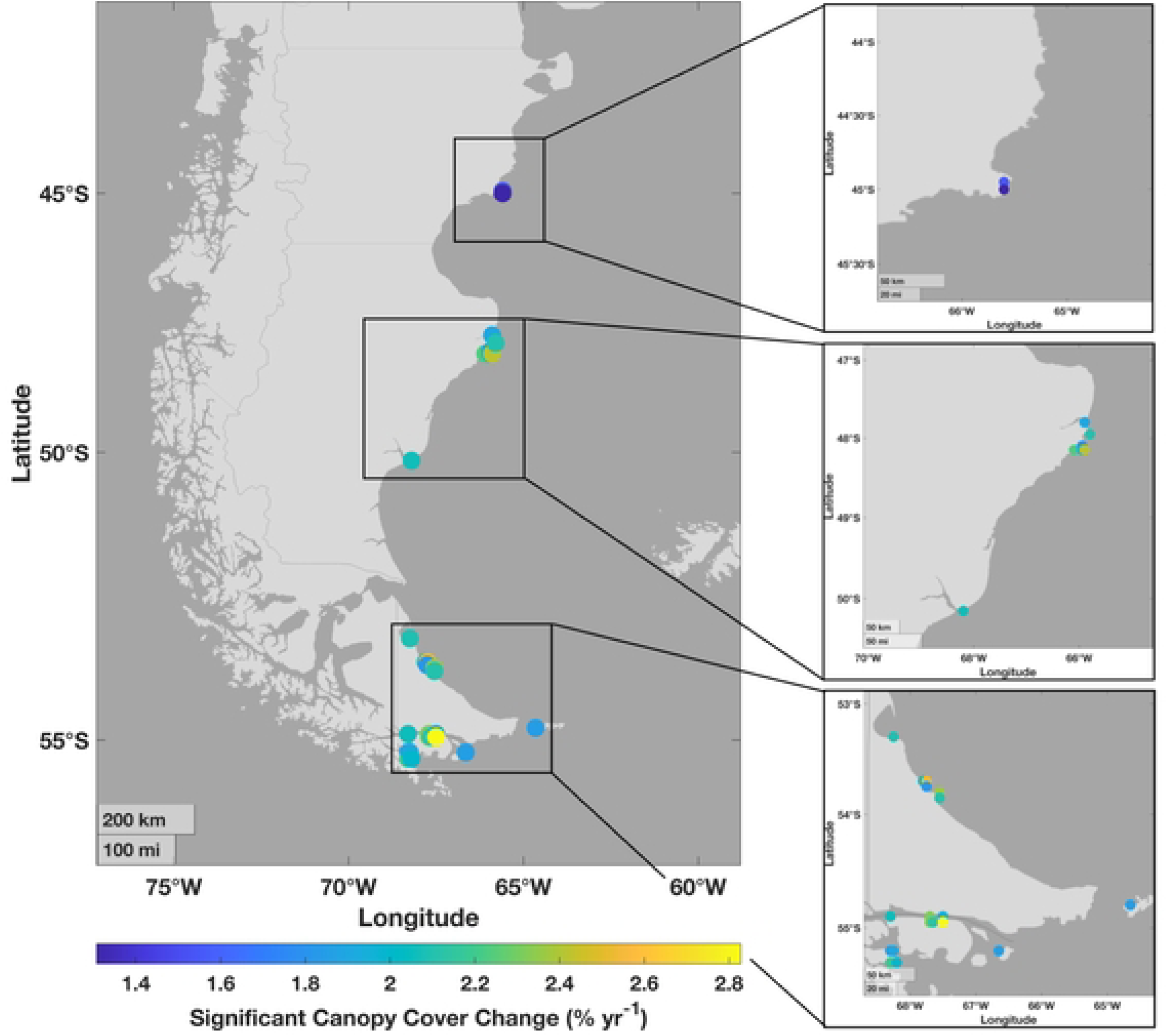
The sites of kelp canopy that displayed a significant change in canopy, (q ≤ 0.05) from 1998-2023 after false discovery rate (FDR) correction. The panels on the right are zoomed in to the three provinces, Chubut, Santa Cruz, and Tierra del Fuego, where kelp canopy is present.

**Fig. 6.**
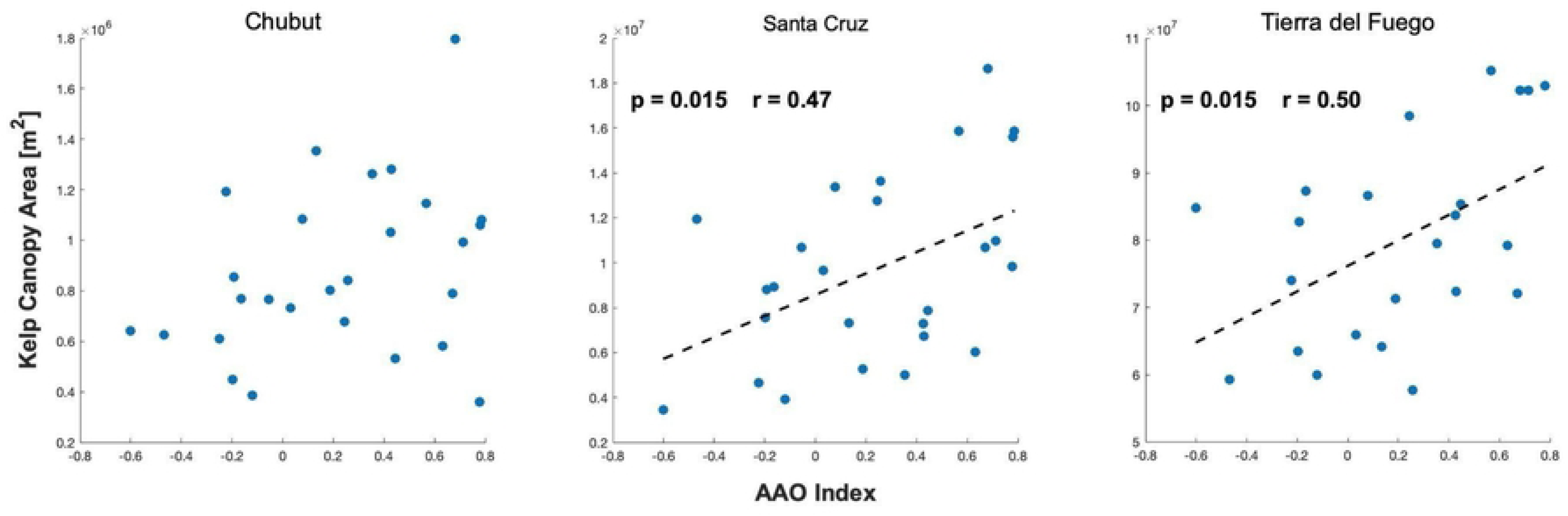
Relationship between kelp canopy area and the Antarctic Oscillation Index (AAO). Black dotted lines in the Santa Cruz and Tierra del Fuego panels show the fitted linear regression (kelp canopy area as a function of AAO).

**Fig. 7.**
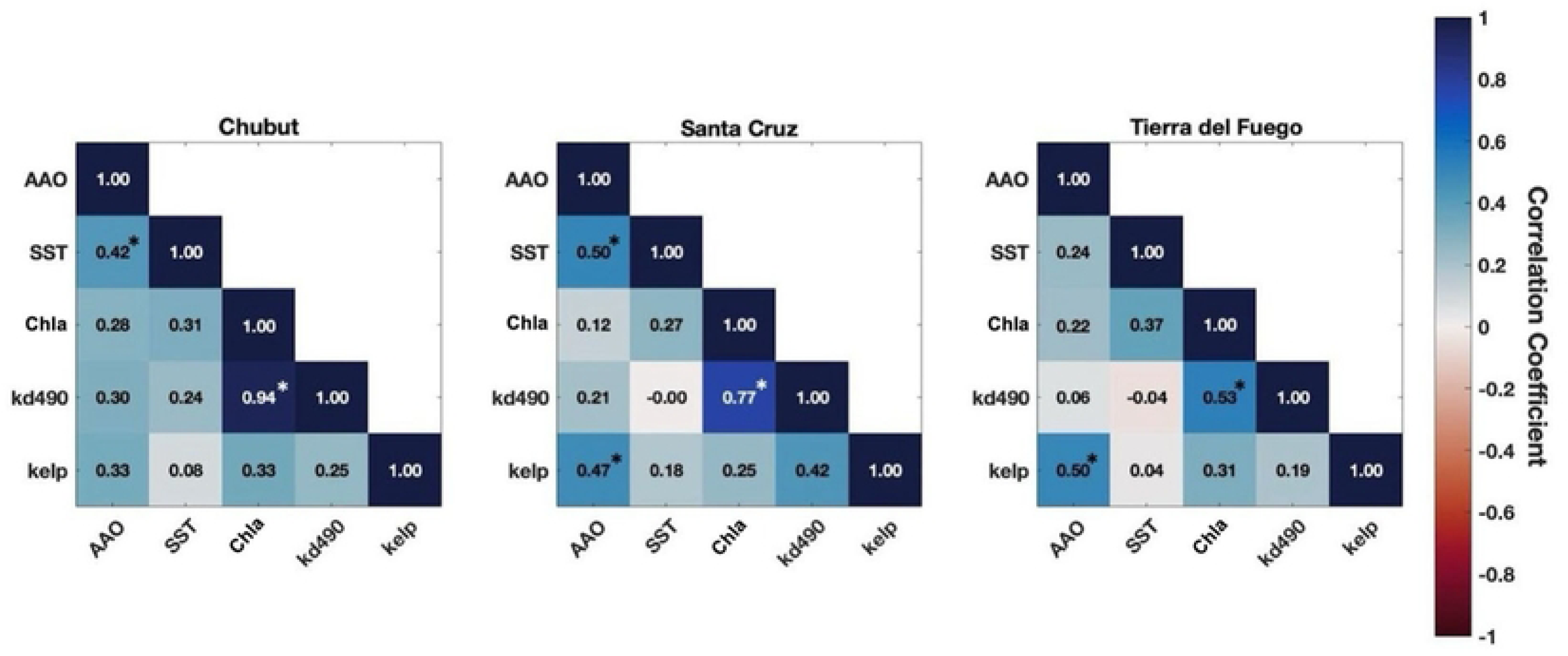
Correlation matrices analyzing the relationships between the AAO, SST, Chla, kd490, and annual kelp canopy area (kelp). Colors represent the correlation coefficient between the variables. Relationships associated with a significant p value (p < 0.05) are marked with an asterisk.

### Light Attenuation and Chla Analyses

Time series of Kd490 were investigated in relation to kelp canopy dynamics in each province, which displayed a non-significant relationship in Chubut (r=0.2505, p=0.2170; Fig. 7) and Tierra del Fuego (r=0.1910, p=0.3826; Fig. 7). However, a relationship between Kd490 and kelp canopy dynamics was observed in Santa Cruz (r=0.4244, p=0.0307; Fig. 7). The relationship between Kd490 and the AAO was also explored but revealed no significant correlation in any province (Chubut: r=0.2988, p=0.1382; Santa Cruz: r=0.2078, p=0.308; Tierra del Fuego: r=0.0619, p=0.7640; Fig. 7). Chla was investigated in relation to kelp canopy dynamics, SST, the AAO, and Kd490 and was not significantly related to kelp canopy dynamics along the coast (Chubut: r =0.3349, p= 0.0945; Santa Cruz: r=0.2456, p=0.2265; Tierra del Fuego: r=0.3089, p=0.1515; Fig. 7). Chla was also not significantly related to SST along the coast (Chubut: r=0.3099, p=0.1234; Santa Cruz: r=0.2677, p=0.1860; Tierra del Fuego: r=0.3691, p=0.0635; Fig. 7), nor was Chla strongly related to the AAO (Chubut: r=0.2841, p=0.1595; Santa Cruz: r=0.1218, p=0.5532; Tierra del Fuego: r=0.2227, p=0.2742; Fig. 7). The relationship between Chla and Kd490 was highly related across all regions—Chubut (r= 0.9396, p< 0.0001), Santa Cruz (r= 0.7690, p< 0.0001), and Tierra del Fuego (r=0.5298, p=0.0054; Fig. 7). This was anticipated because Kd490 and Chla are both sensitive to changes in phytoplankton concentration and both use blue and green band ratios of satellite ocean color observations.

## Discussion

### Distribution and Observed Long-Term Trends of Kelp Canopy

The coastal waters of southern South America contain widespread *M. pyrifera* beds, with some areas displaying persistence and adaptability across many decades [13,21],[34,67]. The results shown here document that the coast of Argentina is home to some of the most extensive *M. pyrifera* forests globally, with approximately 247.61 km² of habitat supporting these kelp forest canopies (Fig. 3). Acknowledged uncertainties in this estimate include fractional cover of kelp-containing pixels (potentially biasing the area estimate high), as well as remote sensing limitations (radiometric or algorithmic sensitivity, near-shore masking, or canopy submergence) that would degrade satellite kelp forest detection (potentially biasing the estimate low). With the exclusion of the Chilean islands that were included as part of Tierra del Fuego analyses, in Argentina alone the analyses found 169.71 km² of kelp canopy area. Of this total area of kelp canopy in Argentina, Chubut has 9.99 km² (∼6%), Santa Cruz has 69.66 km² (∼41%), and Tierra del Fuego has 167.96 km² (∼54%).

This study presents the first regional time series of kelp canopy area in Argentina and shows that these extensive forests are also demonstrating persistence at multiannual time scales. Most individual sites did not show a significant change in annual percent cover on the 5 km scale (Fig. 5). This aligns with previous studies that investigated kelp forest dynamics in Isla de los Estados and the Falkland (Malvinas) Islands that found kelp forests to be stable over time [21,35]. The satellite time series suggested possible positive trends in kelp canopy area at both regional and local scales (Fig. 5). Approximately 7.4% of the kelp sites exhibited significant positive trends over time. Provincially, significant positive trends were observed in 2 of 34 cells (5.9%) in Chubut, 7 of 97 cells (7.2%) in Santa Cruz, and 18 of 236 cells (7.6%) in Tierra del Fuego (Fig. 5). Although significant positive trends were observed across all three provinces, these increases were confined to a relatively small subset of the study area. This spatial variability suggests that local environmental conditions likely influence where kelp forests exhibit significant long-term expansion and highlights the importance of continued monitoring to identify the factors driving these localized changes.

The northernmost range limit of *M. pyrifera* is observed near the northern Chubut border, which aligns with previous research [50,68]. However, potential misclassification in this area may arise due to the presence of the invasive kelp *Undaria pinnatifid*a. *U. pinnatifida* appeared in Puerto Madryn in 1992, and has since expanded its range both within and outside of Golfo Nuevo [69]. Other intertidal algal taxa are *Lessonia,* found in areas of Santa Cruz, and *Durvillea antarctica,* found along the natural protected area of Peninsula Mitre. These species grow in the low intertidal zone and can become exposed during low tide, which could increase the likelihood of misidentification as *M. pyrifera*. However, the methods in this study used a conservative approach to mitigate potential misclassifications; the coarse resolution of Landsat and land masking procedure can miss small and fringing beds but decrease the likelihood of including other taxa [70,71].

While positive trends in the kelp canopy area were detected across the regions (Fig. 5), the lack of early continuous imagery and missing data points can challenge the determination of trends [72,73], and results should be considered with caution until a longer time series can be achieved. Regional differences in these trends are expected due to variations in natural climate oscillations, the period analyzed, and the spatial scale of data. Despite statistical significance, short or fragmented datasets challenge conclusions about long-term changes for systems wherein decadal scale variability is important [72,74]. For example, the results suggest that the *M. pyrifera* area in Argentina is likely modified by low-frequency climate oscillations, which indicates that a 25-year dataset may not yet be adequate to quantify long-term variability with high confidence [72,74].

### Kelp Canopy Response to Environmental Variables

Kelp forests are highly variable across time and space and are subject to a multitude of environmental factors [5,17,75,76]. With multiple dynamics at play, long-term data is critically important in elucidating the factors that are influencing kelp forests habitats. Several known drivers of kelp canopy were investigated, such as SST, seawater nitrate concentration, and turbidity. Seawater nitrate concentrations have been shown to have a strong, negative, and non-linear relationship with seawater temperature in regions such as upwelling systems [32,55,77]. This inverse relationship between seawater nitrate and temperature was also evident in this study region (Fig. S3).

Warm ocean temperatures and low seawater nitrate concentration have been shown to be deleterious factors, especially in upwelling systems in the Northeast Pacific [17,31,78], where nitrate concentrations can be observed to be lower than 1 µmol L^-1^ [31,79]. Along the coast of Argentina, available nitrate was not found to be a limiting factor, which could be explained by the high baseline levels of nitrate in the seawater (Fig. S4). The lowest observed annual mean nitrate concentration was around 7 µmol L^-1^ in Chubut, while the highest levels were near 23 µmol L^-1^ in Tierra del Fuego (Fig. S4). The Chubut province had the lowest nitrate levels and the warmest annual SST (Fig. S4 & Fig. 7). Chubut is closer to the influence of the Brazil current, which is characterized by warm, less nutrient rich waters [35,80]. Conversely, Tierra del Fuego had the lowest annual SST and the highest nitrate concentrations (Fig. S4 & Fig. 7). Tierra del Fuego is more proximal to the FMC, which is a branch of the Antarctic Circumpolar Current [81], and it moves Subantarctic waters to the Patagonian shelf [70]. Kelp canopy area increases south of Chubut toward Santa Cruz and Tierra del Fuego (Fig. 3), and many cells showed significantly increasing kelp canopy area in Tierra del Fuego (Fig. 5). The cold, nutrient-rich waters of the FMC likely contribute to the high biomass and productivity of the more southern region. While a strong relationship between SST and nitrate has been observed, the nitrate data derived from [55] were collected offshore and during one season. Kelp habitats are located inshore and this SST-nitrate relationship may vary along the coast, particularly in areas such as channels and fjords affected by freshwater inputs from glacier melt [82]. Although nitrate was not observed directly, the results observed for Chla—which also did not indicate significant relationships to kelp canopy area—supports nitrate variability not being a primary driver for giant kelp variability, as nitrate is a key driver for phytoplankton growth, similar to kelp. Some cells with negative relationships observed to Kd490 were near coastal cities (Fig. 5), where increased sedimentation can lead to higher turbidity [13].

Kelp forests are light-dependent ecosystems, and the depth distribution, health, and reproduction of kelp is strongly influenced by light availability [5,83,84]. Kd490 is a widely used parameter describing water clarity and turbidity, and increases are associated with higher concentrations of phytoplankton, suspended sediments, or dissolved organic matter, all of which increase light attenuation. The results show that Kd490 was not significantly related to kelp canopy dynamics in Chubut or Tierra del Fuego; however, a relationship was observed at multiple cells in the Santa Cruz province (Fig. 7). The Santa Cruz coast features several coastal inlets, rivers, and bays, including the Deseado, Santa Cruz, and Coig Rivers, as well as San Julián Bay [37]. These river systems transport runoff from agricultural, industrial, and urban activities into coastal waters, influencing estuarine dynamics and reducing water clarity [85]. Increased turbidity is often a stressor for kelp in early life cycle phases via reduced available light at the substrate, which hinders settlement and recruitment of sporophytes [14,86,87]. However, turbidity can also provide nutrients, including nitrogen which is an essential macronutrient for kelp [88]. These added nutrients can enhance primary production and growth, as nitrogen supports increased photosynthetic pigment content, boosting photosynthesis and growth rates [88,89]. Given the high concentrations of seawater nitrate observed along the coast, it is possible that nitrogen availability may enhance photosynthesis and support kelp growth, even under turbid conditions. *M. pyrifera* has been shown to acclimate to local environments through morphological and physiological plasticity [13,90,91], it is possible that kelp in this region may be adapted to persist under nutrient-rich but low-light conditions. Future work can assess the impact of turbidity in the specific, local cells where a relationship between kelp canopy and kd490 was observed.

### Kelp Canopy and Environmental Variables in Relation to Climate Oscillations

The AAO describes the oscillatory pattern in atmospheric pressure between the mid-latitudes of Chile and Argentina and the high latitudes of the Weddell and Bellingshausen Seas. Shifts in the phases of the AAO influence regional winds, SST, and nutrient transport and mixing [92,93]. This study found that the AAO was positively related to kelp canopy area across most of the region (Fig. 6) and was associated with changes in regional oceanographic conditions (Fig. 7). For province-specific results, the correlation to annual canopy area of *M. pyrifera* was found to be significant in Santa Cruz and Tierra del Fuego, but not in Chubut. The AAO was positively associated with SST, therefore negatively associated with seawater nitrate concentration, in Chubut and Santa Cruz, but not Tierra del Fuego. This may indicate a difference in kelp forest drivers between regions, or it may be associated with differences in the temperature-to-nitrate relationship across different latitudes. The differences between the provinces in the relationships observed between annual giant kelp area and the AAO, nonetheless, indicate that the AAO’s influence on kelp canopy is likely stronger in provinces closer to the Antarctic Frontal Zone, where its atmospheric and oceanographic effects are more pronounced. These findings are consistent with other regional studies showing that large-scale climate oscillations, such as the PDO and NPGO in addition to the MEI, AAO, and SAM tested here, can influence kelp forest distribution, abundance, and community structure, as well as key environmental drivers including SST, nitrate availability, wave disturbance, marine heatwaves, and grazing pressure [17,18,26,94].

While relationships between kelp forests and other climate oscillations have been widely studied, the influence of the AAO specifically on Southern Hemisphere kelp forests remains comparatively understudied. The AAO is the dominant mode of atmospheric variability in the southern hemisphere influencing wind patterns, SST, and upwelling intensity across mid-to high- latitude oceans of the southern hemisphere [58]. Variations in the strength and position of the westerly wind belt associated with positive SAM phases have been shown to drive broad-scale changes in ocean circulation, heat transport, and surface mixing throughout the Southern Ocean [95,96]. The SAM and the AAO describe the same large-scale pattern of Southern Hemisphere circulation variability, and thus studies on the SAM provide useful context for interpreting the AAO findings in this study. These AAO related atmosphere patterns may help explain the observed relationship between the AAO and kelp canopy dynamics, particularly in the southern provinces where influence is the strongest. The SAM and the AAO represent the same generalized phenomenon of southern hemisphere circulation variability [41]. The index values of each vary slightly, which is why they were investigated separately in this study. While the SAM was not significantly related to kelp forest dynamics along the coast, the p-values for Santa Cruz (p=0.0693) and Tierra del Fuego (p=0.0561) were marginally significant. A previous study by [74] found that ∼40 years of continuous data is essential for elucidating the patterns of low-frequency decadal oscillations occurring simultaneously as natural variability, especially in highly variable systems such as kelp forests. Since this study spans 25 years, a longer time series could elucidate a relationship between the SAM and kelp forests dynamics further, and will be required to better understand whether the trends assessed herein are robust to decadal scale variability.

## Conclusions

This study presents the first spatially resolved time series of kelp canopy dynamics spanning the coastal waters of Argentina. The long-term record supported by Landsat allows analysis of this dynamic system at broad spatial scales and for time scales approaching those for which trends can be separated out from decadal scale processes. Globally, there are many regions where kelp forests are declining [9,32,94], but the results of this study suggest that kelp forests along the coast of Argentina have remained relatively robust, with some evidence for possible increases in canopy area over the past 25 years. These findings are consistent with previous reports suggesting possible kelp forest refugia in southern South America, and document vast areas of kelp forests in Argentine waters of this region. The scale of the giant kelp forest area identified and the distribution of cells indicating positive kelp forest area trends can inform conservation efforts in this region and others. These findings regarding the AAO being positively related to *M. pyrifera* canopy area across the southern portions of the region demonstrates the importance of climate variability in determining kelp forest status and trends. Despite the abundance of kelp forests in coastal waters of Argentina, the region’s monitoring remains comparatively limited. Overall, these findings suggest strong differences in kelp forest status across different regions, indicating that there are, perhaps, places on earth where kelp forests are resilient. Continuing expansive satellite observing of Earth’s global kelp forest ecosystems will be critical to determine whether these apparent refugia in regions such as Argentina will persist and to gain confidence in interpreting variability in kelp forest area given the likely importance of decadal scale climate patterns.

## Acknowledgments

We thank Julieta Kaminsky and Ted Maksym for providing feedback and engaging in helpful discussions. We also thank the NASA Ocean Biology and Biogeochemistry program (80NSSC21K1429), the NASA Commercial Satellite Data Acquisition (CSDA) program (80NSSC25K0324), and The Nature Conservancy for funding this research.

AA: Data curation, Formal analysis, Investigation, Validation, Writing

CP: Validation, Writing – review & editing

HH: Data curation, Methodology, Writing - review & editing

TB: Conceptualization, Funding acquisition, Data curation, Supervision, Methodology, Writing - review & editing

## Supplemental Figure Captions

Fig. S1. Provincial time series of annual kelp canopy area from 1998-2023. Early data points in the Chubut Province displayed.

Fig. S2. Provincial time series of seasonal (quarterly) kelp canopy area.

Fig. S3. Nitrate data was gathered by data scraping from a previous study by Lara et al., 2010 to establish a regional relationship between temperature and nitrate. A Generalized Additive Model (GAM) was used to analyze the relationship between temperature and seawater nitrate concentration and found the relationship was statistically significant (R²=0.936, p<0.001).

Fig. S4. Annual seawater nitrate concentration (μmol L ^-1^) estimated from sea surface temperature from 1985-2023.

